# Reconciling land sharing and sparing: a conceptual model for land use

**DOI:** 10.1101/2023.11.09.566356

**Authors:** Marco Fioratti Junod, Nicolas Borchers, Brian J. Reid, Ian Sims, Anthony J. Miller

## Abstract

Since the introduction of the terms in the early 2000s, the land sharing versus land sparing dichotomy has sparked considerable discussion stemming from the relationship between food production and ecological function. The theory underpinning these approaches straddles disciplinary and ideological boundaries and is frequently outlined within a general economy versus ecology background. Both strategies have come up with competing models on the relationship linking agricultural production and ecological function.

The present work introduces a conceptual synthesis to unify alternative models under a single coherent framework. A generalized model linking agricultural production with ecological function that can harmonize these conflicting views is presented. In addition, a conceptual shift of paradigm is suggested to approach the issue, moving from yield targets to preservation thresholds.

This article provides the theoretical means to identify the environmental and policy conditions under which land sharing strategies, based on low inputs and low impact per unit of cultivated land, can compete with land sparing approaches for the combined goals of biodiversity preservation and food safety.

## 1 Introduction

Over the last 20 years several disciplines have joined the debate around the trade-off between agriculture and wildlife conservation. This debate is usually framed around the land sparing/land sharing tag, where ‘sparing’ represents intensive high yielding agriculture on a small area of productive land and ‘sharing’ requires wildlife friendly farming with lower yields on a larger area of land. Opinions on the subject reflect a broad range of ideological positions, and tend to cross disciplinary boundaries straddling from economic (Kremen, 2015; J.-M. Salles et al., 2017) to conservation (A. Balmford et al., 2015; Baudron & Giller, 2014) through to developmental studies (Fischer et al., 2017). Recently, the impact of climate change, increased energy costs, population growth and the need for more housing in many countries are adding a new layer of complexity and a sense of urgency to this pivotal debate. While many more trade-offs can be identified around agricultural production, spanning from land ownership and environmental policy enforcement to allocation and distribution of resources, the link between production and preservation of ecological function cannot be circumvented and is the focus of the present work.

The role of conservation and low-intensity agriculture in global land use has often been framed, perhaps starting with the pivotal article by Green et al. (2005) in a debate opposing the two alternative frameworks of land sharing and land sparing. Organic and conservation agriculture, together with other forms of agroecology, are considered as the keystone techniques of the land sharing approach, which is based on the idea that a high level of biodiversity and ecological function can be sustained on agricultural land. By contrast, supporters of the land sparing paradigm argue that aiming for intensive, high-yield cultivation is overall a biodiversity-friendlier approach since it allows production targets to be met using a smaller surface area, therefore freeing large areas for minimally managed natural ecosystems. Occasionally, land sparing is referred to as the Borlaug model, since one of the underpinning objectives of the green revolution and of its chief proponent Norman Borlaug, was to stop the encroachment of agriculture into surviving forests and natural grassland by dramatically increasing production on smaller land areas (B. T. Phalan, 2018). Management of landscape connectivity, which allows otherwise fragmented ecosystems to benefit from ecological corridors favouring long-range migrations, gene flow and rapid recruitment after disturbance, is also a key tenet of the land sparing approach, whereas it is deemed to be largely superfluous under land sharing. While the two approaches are largely alternative in their foundations and landscape-scale application, overlap and blurred borders can occur at a smaller scale, where wild margins and fallow corridors can be part of the toolkit of either approach (Gilroy et al., 2014; Grass et al., 2019; Hodgson et al., 2010).

Most of the field and associated modelling data compiled so far point to an inherent advantage of the land sparing approach (Phalan et al., 2014; Segre et al., 2022), but there are dissenting studies (Steffan-Dewenter et al., 2007) and the issue should not be prematurely considered as settled (Kremen, 2015). Others point to the fact that the optimal choice would be context-dependent, with suggestions that land sharing might be more appropriate in temperate regions and land sparing in tropical climates (Ramankutty & Rhemtulla, 2012). In addition, advocates of land sharing claim that the yield gap between conservation and industrial agriculture is not as large as portrayed, and that in the environmental budget of intensive agriculture there are substantial negative externalities that are not taken into account by most comparisons (Matson & Vitousek, 2006). Some authors also highlight the different ethical frameworks underpinning the two strategies, with considerations other than a purely scientific debate essential to assess the optimal choice (Loconto et al., 2020). Moreover, the dichotomous nature of the debate has also been questioned, with a synthetic assessment of the two philosophies pointing out that the global adoption of either of these techniques would be an undesirable outcome for global biodiversity (Kremen, 2015). The criticism is based on the limits of the reductionist approach that such a binary choice entails, and on the observation that a combination of elements from the two alternatives would offer better perspectives (Baudron et al., 2021).

Whatever the ideological starting point, the trade-off between agricultural production and ecological performance is well-documented and has wide-ranging consequences for global land use and agricultural production. While many additional considerations can be made regarding the social, economic and political aspects of food production, the decision about the optimal level of intensity in the exploitation of each individual parcel of land cannot be avoided. The fundamental question “to what extent can yield be sacrificed for environmental benefits?” lies at the heart of the dilemma. The monetisation of negative and positive externalities – adverse or beneficial effects on the surrounding environment or local communities-is now commonplace in agricultural policies (Pretty et al., 2001). However, if the adoption of practices to reduce negative externalities or increase positive ones results in a yield contraction, and if the production at a landscape scale must meet stable, or even increasing needs, land managers will face a complex trade-off (Li et al., 2020). More specifically, if the improvement in environmental quality following the adoption of the new practices is quantitatively more than compensated for by the loss in environmental quality on the additional land required to be put into production to maintain production targets, the environmental balance would be negative.

## 2. Production and ecological function: trade-offs and trajectories

To provide answers to this conundrum, a conceptual framework is required that links production and ecological function. This is a necessary step, but it is fraught with possible pitfalls when it comes to defining and quantifying the variables involved.

The term production as employed in the current article encompasses both the crop output of agricultural activities and its energetic, mechanical and chemical inputs. While individual inputs are likely characterised by diminishing marginal returns, an assumption of near-linear correlation between aggregate inputs and yield is made here (J. M. Salles et al., 2017). The assessment of intensity using output as a proxy has been a staple of land sharing/sparing studies. The explicit negation of a linear linkage between the two will be nevertheless taken into account when examining different conceptual approaches. In this case a distinction between inputs, here termed intensity, and outputs, designated as yield, will be introduced.

As for ecological function, biodiversity and other ecosystem services have been historically favoured as proxies to assess the response of an environment to changes in land use. However, the choice of indicator remains controversial. In most studies, biodiversity is approached at a landscape scale and with a focus on above-ground taxa, which benefits from solid foundations using surveys and structured indexing (Clergue et al., 2009). However, the observable biodiversity is often a partial and imperfect measure of the ecosystem at large and distinct from the ecological function a landscape delivers (Hagan et al., 2021). Similarly, the very concept of ecosystem services have been the object of intense scrutiny in recent years, with criticism focusing on their alleged anthropocentrism and utilitarianism (Muradian & Gómez-Baggethun, 2021; Schröter et al., 2014) as well as the widely divergent scope of its application in different contexts (Kull et al., 2015). Nevertheless, the use of biodiversity indices and ecologically-relevant elements of the ecosystem service framework are still pivotal to our understanding of global patterns in response to anthropic disturbance. Our proposed hybrid concept of ecological function is meant to encompass a variety of performance indicators, taken individually or as bundles of largely collinear metrics. As an illustration, to test the proposed models below we used alpha diversity metrics for above ground taxa (B. Phalan et al., 2011). Given a degraded reference point however, beta-diversity dissimilarities would be an equally acceptable choice (Bray & Curtis, 1957).

A broad and all-encompassing definition of these variables-production, intensity and ecological function-makes it possible to create a common scaffolding to frame a variety of published models. These proposed models link what was here broadly categorised as ecological function versus production. We here propose a five-way taxonomy using linear relations or simple curves.

First, there are the convex and concave models (Figure 1 (A)), usually presented in parallel and described by curves borrowed from species-density or survivorship functions (Green et al., 2005; B. Phalan et al., 2011; J.-M. Salles et al., 2017). The convex model (Green et al. (2005) use convex and concave as seen from the origin instead) involves a mild decline in ecological function at low production followed by a sharp downturn at high production. Its mirrored counterpart across the diagonal, the concave model, is characterised by a steep downward trend at low production, followed by an anelastic phase. When it comes to sorting the trade-off between production and ecological function the optimal production values lie at the lower end for the convex model and at the higher end for the concave model. The third conceptual model (Figure 1A)) is rarely expressed in mathematical terms, but the formula underpinning it is implicit by its theoretical definition. Some authors insist on the drivers of the land use debate being chiefly ethical in nature (Loconto et al., 2020). A radical interpretation of this approach, denying inherent conservation advantages of any specific production level other than in an ethical dimension, would result in what we will call the “ethical” model, in reference to its theoretical foundation rather than its perceived superiority. The only points of the plane satisfying this relation are arranged in a downwards linear trend along the diagonal (Figure 1 A).

**Figure 1.**
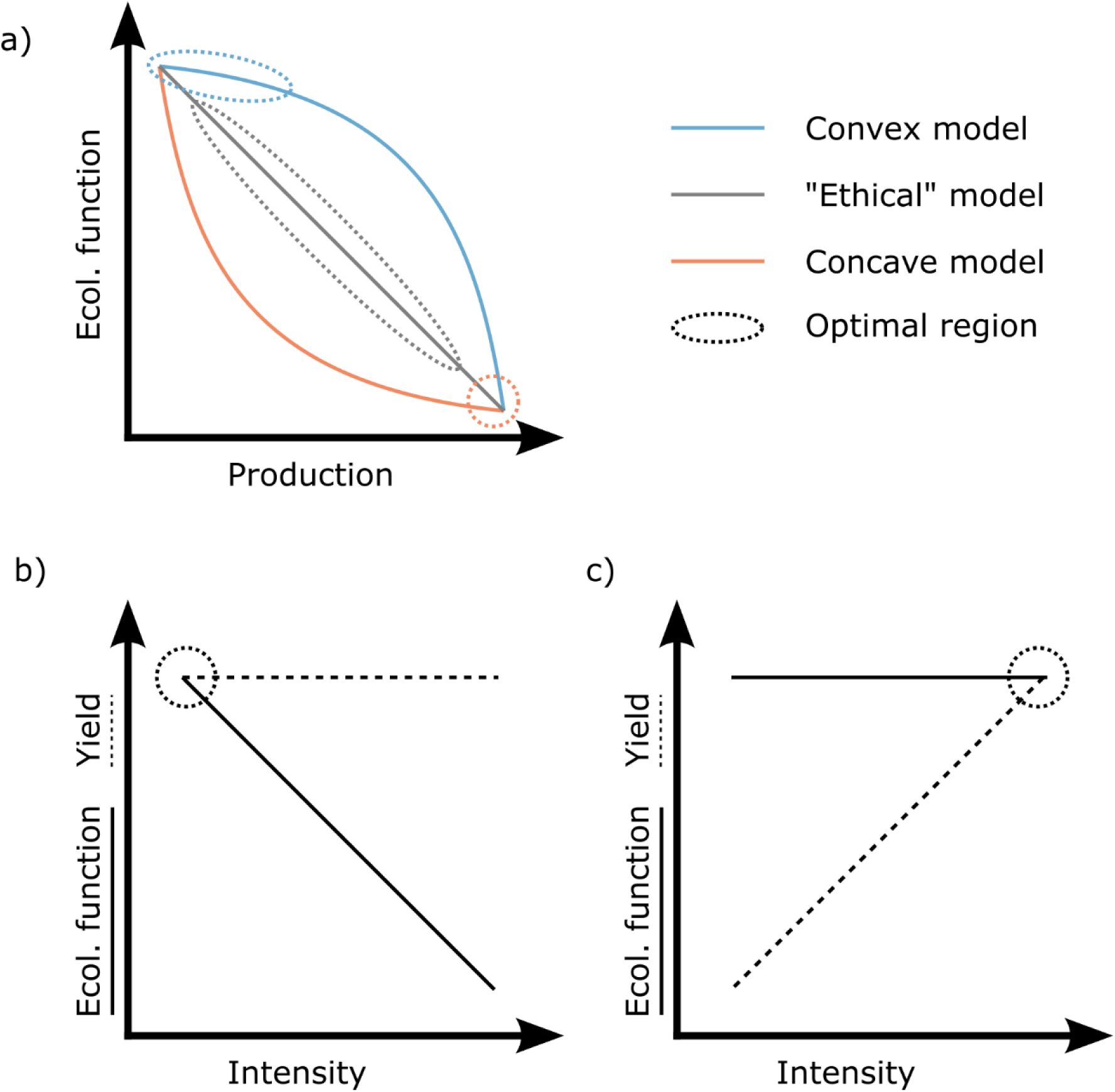
(A) a graphical summary of predictions for the convex, concave and “ethical” models. The oval shapes highlight the advantageous intensity/function configurations. (B) The no yield trade-off model. (C) The no function trade-off model.

The remaining two models are characterised by their rejection of one of the two implicit trade-offs accepted by the previous ones, which means that the production variable will be split between the now diverging components of yield and intensity. The no yield trade-off model (Figure 1 (B)), while accepting the reduction of ecological function along an intensity gradient, as in the ethical model, postulates that the same levels of production reached by high-input systems can be obtained by conservation-oriented systems. While most proponents of de-intensification recognize a yield gap with conventional high-input agriculture (Wilbois & Schmidt, 2019), some authors outright reject the claim that agricultural output is input-limited (Altieri, 1999; Vandermeer & Perfecto, 2005). This view is fundamentally a variation of the ethical model, that introduces a decoupling of production into yield and intensity components, and involves the presence of an optimal point at high yield and low intensity. Finally, the no function trade-off model (Figure 1(C)) is based on exactly the opposite premise: a direct correlation between intensity and yield is accepted, but the reduction of ecological function at high production is rejected, making intensive agricultural systems always advantageous. This model is interesting as a theoretical exercise but might be applicable only in specific real-world circumstances, for example where agricultural production occurs in extremely poor or degraded environments.

## 3. A unifying approach: the sigmoid response model

Except for the speculative frameworks that deny the two fundamental trade-offs between yield and intensity and between intensity and ecological function (Figure 1 B & C), which better belong in the theory rather than the practice of land use (De Ponti et al., 2012; Dudley & Alexander, 2017; Kravchenko et al., 2017; Outhwaite et al., 2022), the models describe to differing extents patterns that can be observed in real agroecosystems, albeit with the simplifications required at high levels of generalisations. The convex model describes well the resilience of natural ecosystems to small levels of disturbance (Figure 1A). The concave one illustrates the relatively inflexible response of ecological function under marginally increasing levels of stress. The ethical model fits the cases where transitions in ecological functions occur near-linearly at mid-range agricultural production (Figure 1A). While these three models are conceived as mutually exclusive – and they are in their original formulation – we argue they collectively describe a single complex response pattern. This response pattern is better approached and described starting from the well-researched theoretical framework of alternative stable states (Beisner et al., 2003; May, 1977). As observed in many seminatural environments, the response to stress is not a linear one but tends to coalesce around areas of higher stability. In the same way, while agricultural intensity can be seen as a continuous gradient, the environmental response to it is in most conditions segmental and almost discrete (Seppelt et al., 2016).

At low production, biological buffering systems can compensate for exogenous stress and the resilience phase is maintained (Phelan, 2004). Total biodiversity is not affected until the tolerance threshold of a sizeable minority of species is reached. As for ecosystem services, functional redundancy ensures that the loss of some less resistant taxa does not interfere with general trophic chain stability and biogeochemical cycles (Jurburg & Salles, 2015). With increasing production, a phase of transition follows, where the loss of key taxa has a cascading effect on the provision and catalysis of environmental and ecological services. According to the nature of the system and the measured parameter, this phase can be either a sudden collapse or a more gradual decline with resilience and buffering still working to an extent, but not enough to stem the decline from the original threshold. With increasing stress from agricultural activities, a new stable state is reached, the tolerance phase, with its own mechanisms of resilience and buffering (Cropp & Gabric, 2019). A simplified set of functions is performed by assemblages of organisms largely tolerant to stress. Specific elements are depleted until they reach the threshold of their recalcitrant pools. It is possible to imagine this state as the resilience phase preceding a new transition. Indeed, multiple stable states have been described in natural ecosystems subject to disturbance (Moffett et al., 2015). The presence of more than two alternative stable states in agricultural systems is more unlikely, but it should not be discounted. The model we are proposing in its basic form does not cover multiple stable states with stepwise transitions, but it could still be used to cover discrete parts of the intensity range.

For the formulation of a model suitable to represent the transition between alternative stable states in an agricultural context, we used an implementation of the sigmoidal response (1), a common stress response function in many biotic systems. The P independent variable can be taken to represent production, expressed as a fraction of the maximum attainable level, which is assigned a unit value. *f_max_* is taken to be the level of the ecological function under investigation at production zero, before stress is applied. *f_c_* refers to the core function, i.e. the surviving level of ecological function in the simplified stable state, expressed as a fraction of *f_max_*. *σ* (sensitivity) is a parameter determining the slope of the function in the transition phase. Higher values are associated with a steeper transition. While no assumptions are made about hysteresis, and the sigmoid response model is not meant to describe a time series, it is worth noting that steeper transition phases are most often associated with irreversible change (Meyer, 2016). *E* is the ecological efficiency of the system, defined by the value of P at which the function reaches the mid-point of its transition phase (Figure 2).

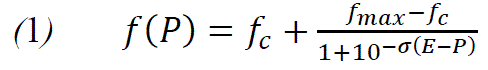

**Figure 2.**
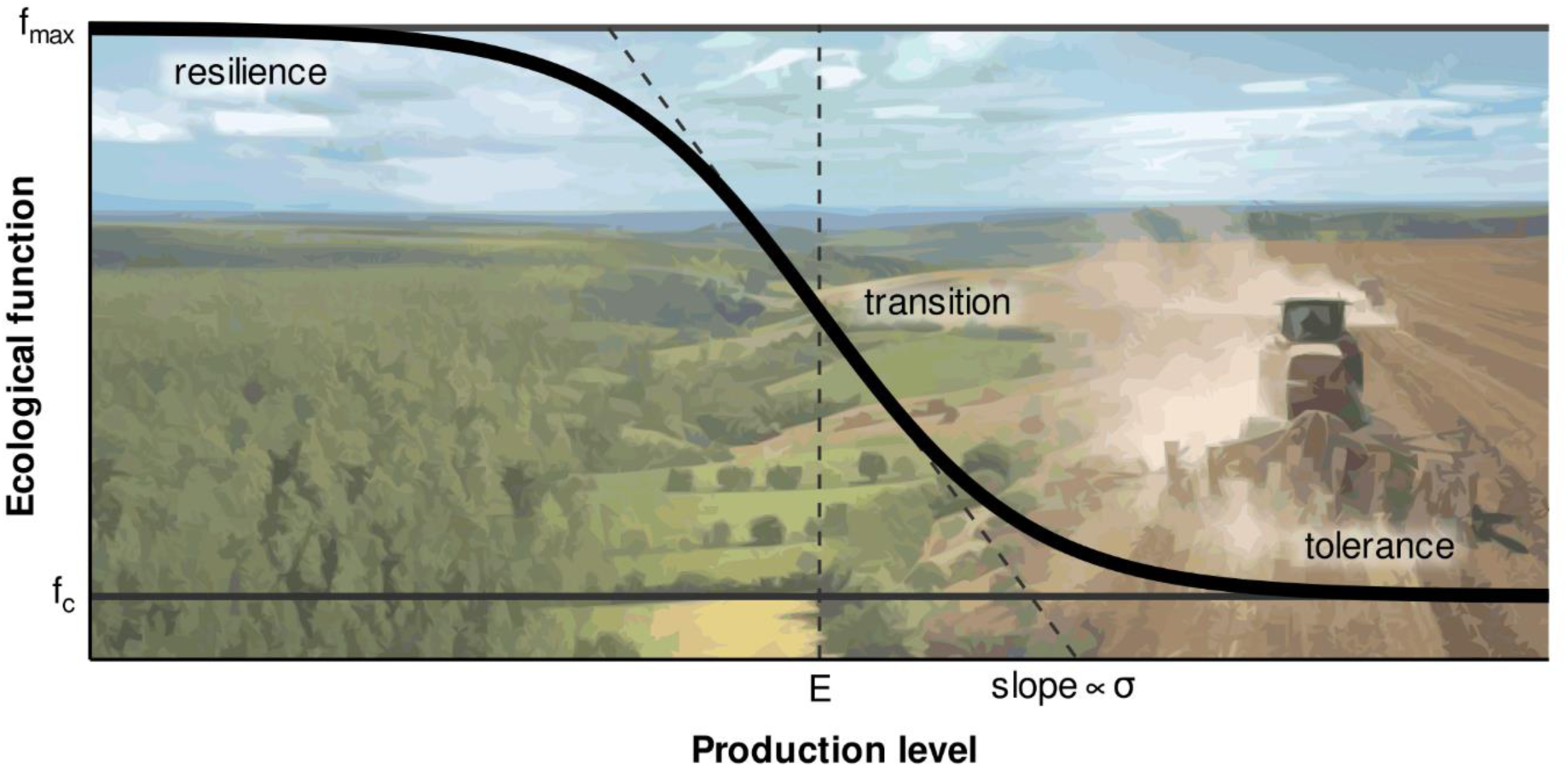
Parameters and component phases of the sigmoidal model.

The function in Equation (1), in addition to the central case of a clearly identifiable sigmoidal shape with a linear transition phase connecting the two asymptotic phases, resilience and tolerance, is flexible enough to approximate the other models discussed previously above (Figure 3). The convex model can be generated using a high value of E and a low value of *σ*. Conversely, the concave model is obtained with values of ecological efficiency approaching 0 and medium *σ* values. Even the linear ethical model can be approximated with a central value of 0.5 for ecological efficiency and an extremely low sensitivity to expand the transition phase. The two no trade-off models are similarly approached, and their graphic representation looks the same when P is taken to represent yield. However, they can be separated theoretically by using an extremely high ecological efficiency in the case of the no yield trade-off model and a *f_c_* value equal to *f_max_* for the no function trade-off.

**Figure 3.**
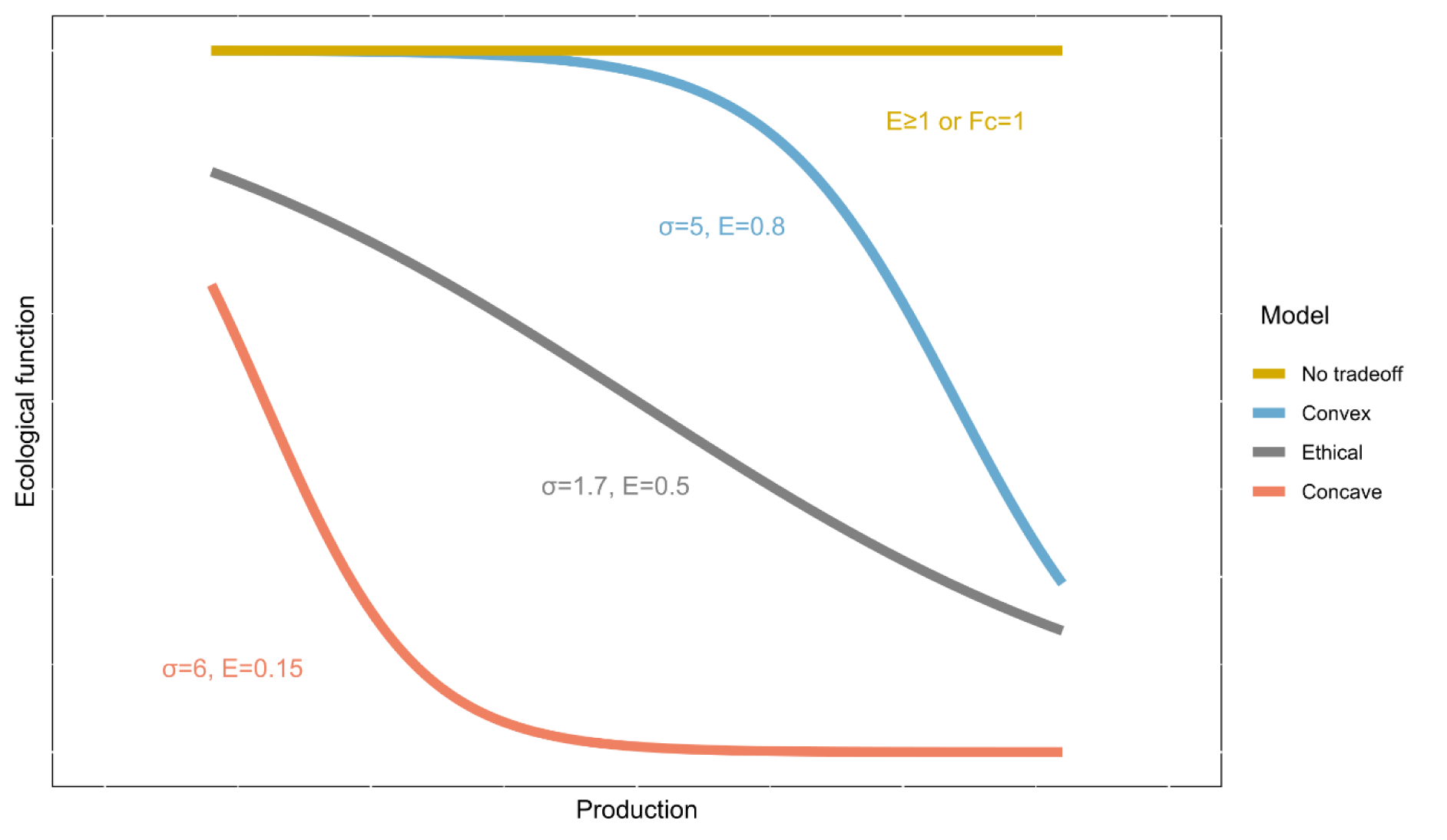
Approximation of the models presented in the proposed taxonomy with the logistic sigmoid equation.

A conceptual representation of the simplified models along these lines is also helpful to understand the mechanisms at play that make one of the phases of the sigmoid curve dominant over the others, generating the impression of a simpler curve. Convex type relations involve a high level of resilience driven by substantial functional redundancy and are followed by a rapid collapse to levels that are just a tiny fraction of the extensive diversity originally present. Concave curves are observed when the initial level of functional redundancy is quite limited and where the core function level is proportionally higher. The ethical model assumes the absence of stable states, at least within the considered range of stress levels. Even the extreme no-trade off models, for yield and function, become clearer to grasp when they are interpreted respectively as the ability to carry out production entirely within the resilience phase of the system and as production occurring in a system that is already in a simplified and depleted state from the start.

Summarising, we argue that all commonly theorised production/function models are just special cases of a more universal sigmoidal relation, where the dominance of one or two of the three phases masks the presence of the others (Cormont et al., 2016). Three-phased configurations of the sigmoid response are actually the norm for a variety of parameters. Density functions for single species offer a good insight into the mechanistic reasons behind the sigmoidal nature of community response to agricultural intensification and the configurations where one or more phases of the sigmoid are emphasised. Phalan et al. (2011), based on comprehensive databases of tree and bird species from Ghana and India categorised the response of individual species to a gradient of agricultural intensification into four main categories, each characterised by a typical yield-density curve type (A. Balmford et al., 2015). The majority of surveyed species could be clearly assigned to one of these archetypal clusters (Figure 4 B). On one side are “losers”, species that show a declining abundance along an intensification gradient, and that form the majority of surveyed species in agricultural context. These can be further divided into two clusters. Species experiencing a population crash at low intensity have a typically concave density curve. At landscape level these species tend to favour land sparing settings. On the opposite, species that tend to preserve good densities at all but the highest levels of agricultural intensification can have their density response represented by a convex curve and fare better in land sharing contexts. The other side is made up of “winners”, a small number of species that benefit from agricultural conversion, either through niche creation or reduction of competition. This second group also comes in two flavours, mirroring the phenomenon described for “losers”. On one hand some species show a convex density response, with a sharp increase in numbers occurring at low levels of intensification and plateauing at higher level. Other species remain at low densities for all but the highest levels of the intensification gradient and show a characteristically convex growth profile.

**Figure 4.**
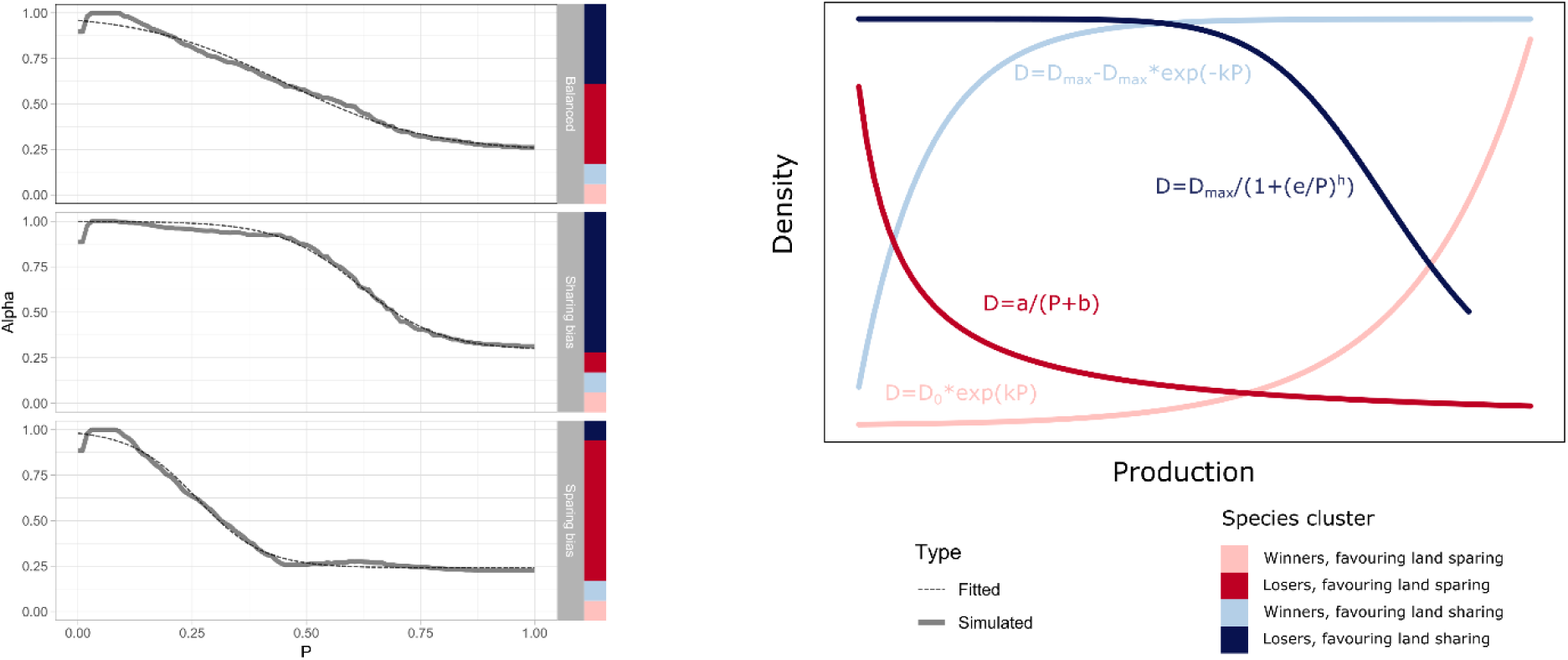
(A) shows a species richness curve generated with simulated communities of a thousand species each. On the x axis is production (P) and on the y axis is the number of species recorded for each production level as a fraction of the highest level within the production continuum. Species are counted if present with 10 or more individuals at any given production level. The simulated data is shown with the full lines, whereas the corresponding fitted sigmoid response curve is shown with a dashed line. The proportion of species allocated to species cluster is shown by the vertical stacked bars to the right. All species were assigned to one of four species clusters, whose baseline density curves are shown in Figure 4 (B). These density curve types are adapted from Phalan et al. (2011) and describe the typical response of species to agricultural intensification based on tree and bird species data collected in Ghana and India. In the simulation, each species allocated to a species cluster type is assigned to one of a hundred curve random variations within the same family. For the “winners, favouring land sparing” cluster, exponential growth curves were adopted with random k values from 3 to 6 and corresponding a values constrained so that for P=1 the value of D was equal to 100. For the “winners, favouring land sharing” cluster, a curve from the exponential plateau family was randomly allocated, having D*_max_* set to 100 and k comprised between 4 and 10. For the “losers, favouring land sparing”, a hyperbolic function was adopted with a random value of a between 1 and 5 and the corresponding values of b set so that the function has a value of 100 when P is 0. For the “losers, favouring land sharing” cluster, a Hill’s decay function with D*_max_* set to 100 and random values of e between 0.4 and 0.6 and of h between –2 and –20.

**Figure 5.**
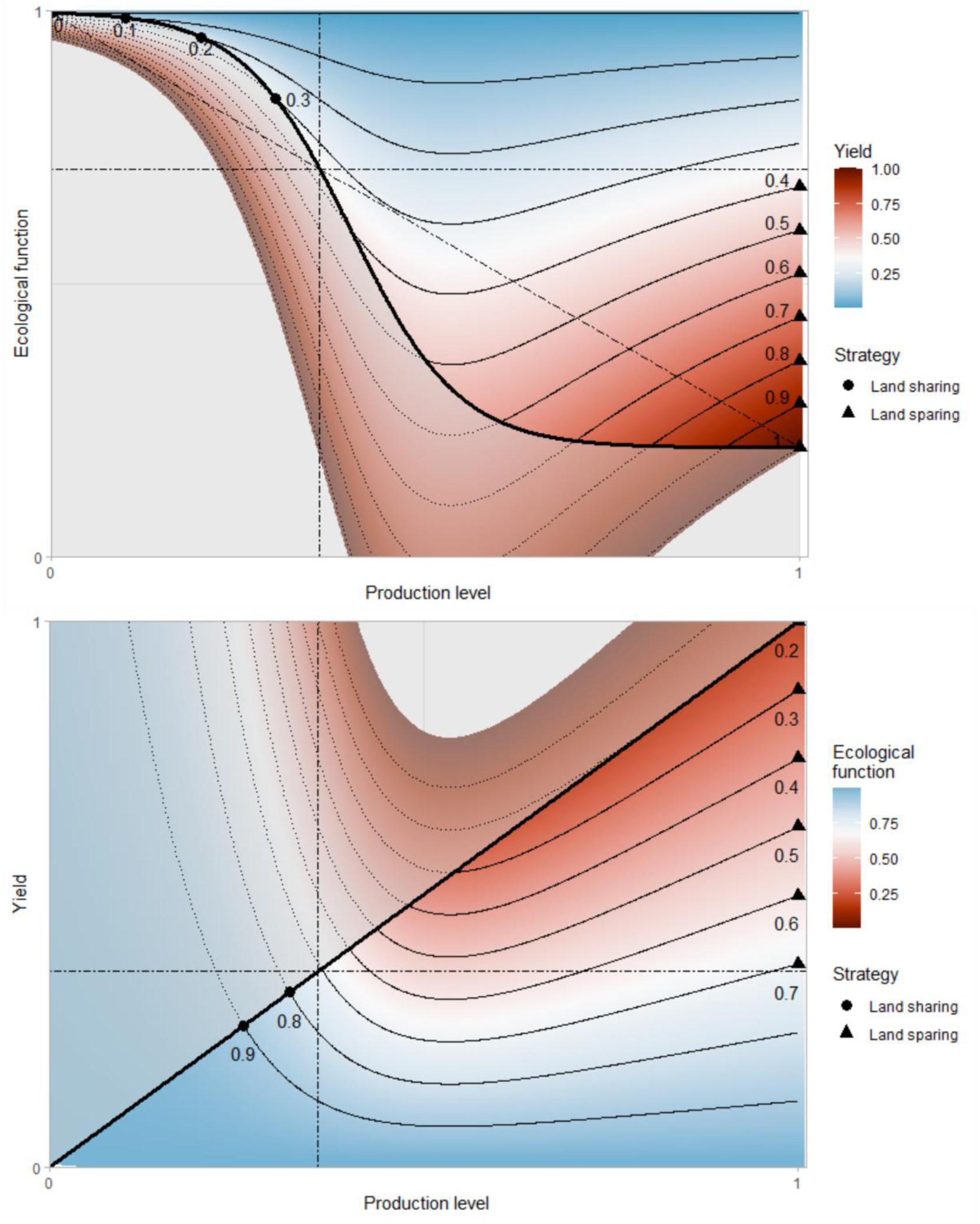
(A) shows a sigmoid with an F*_c_* value of 0.2, a sigma of 6 and an E value of 0.4 as an envelope to isokarps. The background colour and corresponding isokarps indicate the yield gradient. The points show the level of production optimal for each reference yield. The dotted-dashed line shows the neutral yield level, ie the production target at which land sharing and land sparing solutions are equivalent. Above this threshold, land sparing (P=1) is advantageous, below this threshold, the optimal production is equal to the target yield and falls in the land sharing region. Figure 5 (B) shows the isoerg envelope for the same sigmoid curve. The background colour and corresponding isoergs indicate the gradient of ecological function. The points show the level of production optimal for each ecological function level. The dotted-dashed line shows the neutral ecological function level, ie the value at which land sharing and land sparing solutions are equivalent. Below this threshold, land sparing (P=1) is advantageous, above this threshold, the optimal production falls in the land sharing region

While none of these individual species’ clusters show a typically sigmoidal, three-phased response to intensification, it is possible to show that communities made up of combinations of these groups in proportions registered in nature can generate an aggregate species richness profile very accurately described by the proposed sigmoidal model (Figure 3 A). A simulation run on mock communities, each comprising a thousand species belonging to the four different categories, show highly distinctive species richness profiles. With the percentage of “winners” kept to one fifth of the total – a quote consistent with surveys (B. Phalan et al., 2011) – the ratio between “convex” and “concave” winners is shown to dramatically change the overall shape of the function, shifting the weight towards specific phases of the sigmoidal response. An even partition between the two results in the linear, transition phase of the sigmoid to almost completely obscures the asymptotic phases, approaching the resulting function of the previously described ethical model. A prevalence of “convex losers” results in a strongly elongated resilience phase, with higher productions achievable without substantial diversity loss. Conversely, a prevalence of “concave losers” results in a shortening of the resilience phase and a parallel elongation of the tolerance phase of the sigmoid, with wide scope for intensification at high levels without further decreases in diversity.

In all the above-mentioned cases, fitting the proposed sigmoidal function to the simulated data results in very high coefficients of determination (R^2^>0.99), confirming the mechanistic basis for the theoretical foundation of the model. The same observed pattern could also explain why landscape scale ecosystem services based on the direct or indirect activity of biotic communities have also been associated with a sigmoidal development in response to disturbance (Locatelli et al., 2017; Tscharntke et al., 2012). Along the same lines, transition between steady states has been used to interpret long-term trends like soil carbon content following changes in management intensity (Janzen et al., 1998). Even geochemical functions that depend on the activity of biotic communities, such as soil nitrous oxide emissions in response to increased production, have been described with a characteristic shape associated with steady states (Hickman et al., 2017). These steady states, with asymptotic phases linked by a linear transition, are probably driven by a saturation of the processing capabilities of core microbial communities under high fertilisation.

## 4. A shift of paradigm: production-first to preservation-first

Despite structural criticism about the inherent crude reductionism of the approach, land sharing and land sparing as extremes of a conceptual continuum are still pivotal concepts in the formulation of global land use strategies. The lexical and ideological nature of the debate can be rejected, but the decision of an optimal level of production for individual agroecosystems is inescapable and the decision to adopt extensive or intensive farming practices on a specific extent of land brings back the dualism in full force. Identifying which of the two clusters of management techniques has structural advantages in specific conditions is a valuable contribution for land management policies (Phalan, 2018). In particular, since the adoption of a single approach at a global scale is deemed an undesirable option (Kremen, 2015), instruments for allocating land in specific circumstances are required. Gaining a foothold in the understanding of general patterns linking agricultural production to biodiversity and ecological function is a necessary step in this direction.

The level of production, P, is defined here as constrained between 0, on non-farmed land, and 1, for the intensity generating the maximum attainable yield. For a given unit of land, we define s as the ratio of land under productive use and 1-s the unfarmed, wild fraction. The yield of the plot (Y) will be therefore

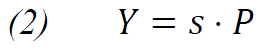

Similarly, the ecological function of land under agriculture at a productivity P can be represented by a function f(P). It is convenient to define f(P) as a spatial density of ecological function, so that the aggregated ecological function for a unit of land combining cultivated and uncultivated fractions can be expressed as

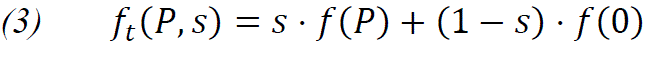

This model is not explicitly taking into account scale effects in the preservation of ecological function across the landscape. Such effects obviously exist and often take the shape of positive feedbacks in ecosystem service dynamics (Segre et al., 2022) or reduced performance due to fragmentation and minimum habitat requirements (Andersson & Bodin, 2014) but are beyond the scope of this streamlined conceptual model. A dynamic definition of f(P) compensating for landscape cohesion would moreover result in more accurate predictions (Villard & Metzger, 2014). Similarly, in the context of broad conceptualisation, no effort was made to integrate a possible divergence between yield and economic values of commodities (Sidemo-Holm et al., 2021).

Substituting (2), we obtain

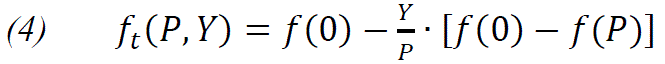

When keeping Y constant, equation (4) can represent the variation of ecological function at constant yield along a sharing-sparing continuum. The resulting curves (Figure 4 (A)) can be named isokarps, from Ancient Greek ἴσος (equal) and κᾰρπός (harvest). These curves can be plotted on an ecological function versus productivity plot. Since

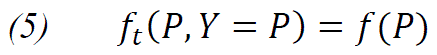

the constraint on permissible values of P draws the envelope of f(P). Isokarps for P values left of this curve are not permissible given the constraint of a limited surface, and are represented by dashed curves. Furthermore, points representing authentic land sharing scenarios are organised along the f(P) curve, while points representing land sparing scenarios are located along the vertical axis (P=1). The area between these two limiting curves represents intermediate solutions between sparing and sharing.

Using the same mathematical scaffolding already introduced, it is possible to rearrange (4) to express the yield as a function of productivity for a given ecological function:

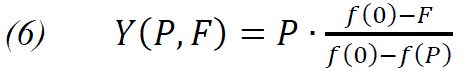

(6) therefore represents curves of constant ecological function, which we will call isoergs, from Ancient Greek ἴσος (equal) and ἔργον (action, function). These can be plotted on a yield versus productivity plot (Figure 4(B)). The line Y=P delimits the permissible values of P and is the equivalent of the f (P) curve on the isokarp plot. Isoergs for P values left of this line are not permissible given the constraint on minimum production on a limited surface, and are represented in dashed curves. By analogy with the isokarp plot, points representing land sharing scenarios are organised along the Y=P line, points representing land sparing scenarios are located along the vertical axis (P=1), and the area between these two limiting lines represents intermediate solutions between sparing and sharing

Both approaches are equally useful to represent respectively ecological function as a result of production and yield, or yield as a result of production and ecological function. Traditional, production-first, land use models are conceptually based on the constraint of meeting a defined production target. The isokarp space can therefore turn from a mere representation tool into a conceptual space where ecological function is maximised following the introduction of enforced agricultural production targets. However, we argue that such a production-first approach is conceptually unfit to correctly frame the land-sharing issues. The concept of yield targets in a land use context is improperly borrowed from the jargon of former planned economies (Dovring, 1982; Xianling et al., 2016). Such targets are not grounded in the current economic reality of land management, where profit is always maximised by rational players and only constrained by externally enforced limits or positive incentives. Externally enforced limits, in the context of the land sharing / sparing debate, are bound to take the form of preservation constraints. We therefore introduced the concept of preservation thresholds (*θ*) to replace yield targets to orient decisions about land management. This change of paradigm involves switching away from the setting of a target for food production and then maximising ecological function within this limit, which might not be sustainable for either humans or the natural ecosystems. We advocate instead assessing a minimum limit of ecological function necessary for ecosystem and human survival and second economical behaviour by maximising production within this constraint. This change can be conceptualised using isoergs. When *θ* replaces F in (6) we enter a conceptual space where yield is maximised within the constrained of an enforced preservation target. The resulting isoerg space becomes a precious tool to address the land sharing / sparing debate from more realistic premises.

## 5. Synthesis: approaching the land sharing / sparing debate

We argued earlier that realistic ecological functions are in general best represented using a sigmoid shape. We introduced above a typical sigmoid function based on the logistic function (equation 1), but the following will be valid for any sigmoid function, which can be defined in this context as any monotonic, strictly decreasing function which is at first convex, then concave.

When f(P) takes the form of a sigmoid, it can be shown that the isokarps f*_t_* (P,Y) are necessarily unimodal with one minimum in respect to P on the interval 0 ≤ P ≤ 1. It follows that for each given yield, optimal solutions i.e. scenarios that maximise average ecological function on the whole plot of land can only be found on either end of the sharing-sparing continuum (Figure 4 (A)). Specifically, the transition point is located where the chord connecting the sigmoid at P=0 and at P=1 intersects the same sigmoid. This chord can be termed the critical chord by analogy with Green et al. (2005). To the right of this point, optimal production levels fall in the land sparing region and require P=1. To the left of this point, optimal production levels fall in the land sparing region for values of P.

It is however useful to move to the isoerg-based conceptual space to explore the relation between yield and preservation thresholds (Figure 4 (B)). The level of P identified by the intersection of the sigmoid with its critical chord allows to determine the value of *θ* at which land sharing and land sparing strategies do not have inherent advantages. For lower values of preservation threshold, land sparing at P=1 will have a competitive advantage, whereas for higher values the optimal P levels will fall in the land sharing region. Except when θ is lower than f*_c_* – i.e. the preservation threshold is never overshot and land sparing always advantageous when aiming at maximal possible yield – it is possible to identify the peaks of each isoerg along the diagonal as the land sharing optima, and the ends of the isoergs along the P=1 axis as the land sparing optima respectively. The relative height of these two points determine which is advantageous for maximising yield, for given combinations of the three parameters of the sigmoid function, and for each given level of preservation threshold.

## 6. Discussion

The assumption of a sigmoidal link between production and ecological function, and the existence of alternate stable states underpinning it, is an effective conceptual framework to interpret the effect of various land management options. In particular, it can explain both the successes and possible malfunctions of conservation and regenerative agriculture practices. More specifically, the model can be used to predict under what conditions such practices are more likely to yield significant benefits, and in what conditions their use is likely to incur heavy trade-offs. Fitting a sigmoidal response curve can be achieved when few datapoints are available, linking production levels to observed biodiversity of ecosystem function within the same landscape and scale. In this way, many of the observed instances of failure of de-intensification programmes to maintain profitable yields find a convincing explanation, even when they succeed in improving biodiversity and ecological functions (Leifeld, 2014). When agricultural systems operate at the high-intensity end of production, well into the tolerance phase of the system, small gains in ecological function will normally come at the price of a significant reduction in yield (Seufert et al., 2012). The typically diminished response to increasing levels of disturbance in the degraded stable state of intensively cultivated arable land plays strongly against the viability of attempts at de-intensification (Millard et al., 2021). However, this does not mean that there is not a place for species-rich rotations, cover crops, conservation tillage and organic amendments in modern agriculture. Indeed, the same sigmoidal response framework offers clear indications as to the conditions where they can express their full potential. To identify them, we can explore the ways the equilibrium of a system can be shifted in favour of land sharing. Among the four variables considered so far, the three parameters of the sigmoidal response curve and the preservation threshold, two (the core function and sensitivity) are specific to the ecological function and the environmental context, and cannot be manipulated. They can, however, orient choices in land management. According to the isoerg conceptual model, raising the core function does not influence the production level at which sharing and sparing are equivalent (Figure 6), but it raises the preservation threshold at which the two strategies are equivalent. When less of the original ecological function is lost at high production, higher enforced protection is required to disincentivise maximum production. De-intensification techniques and extensive cultivation strategies are anticipated to suit better settings where the difference between the pristine state and the post-perturbation state is more pronounced (Standish et al., 2014), which is to say lower core function values. Identification of such areas is therefore key. On one hand, it is possible to argue that complex and rich systems – like the ones in pristine tropical areas – are able to retain substantial residual function even after disturbance. On the other hand, in poorer and simplified starting contexts, like the ones likely to be encountered in historically cultivated temperate areas, the subset functions that have survived centuries of anthropic activities can be thought to be less impacted by additional stress, making the difference between f*_max_* and f*_c_* less pronounced. Whichever of the two patterns prevails for the ecological function under consideration is likely to have a big influence on the prospects of success of land sharing.

**Figure 6.**
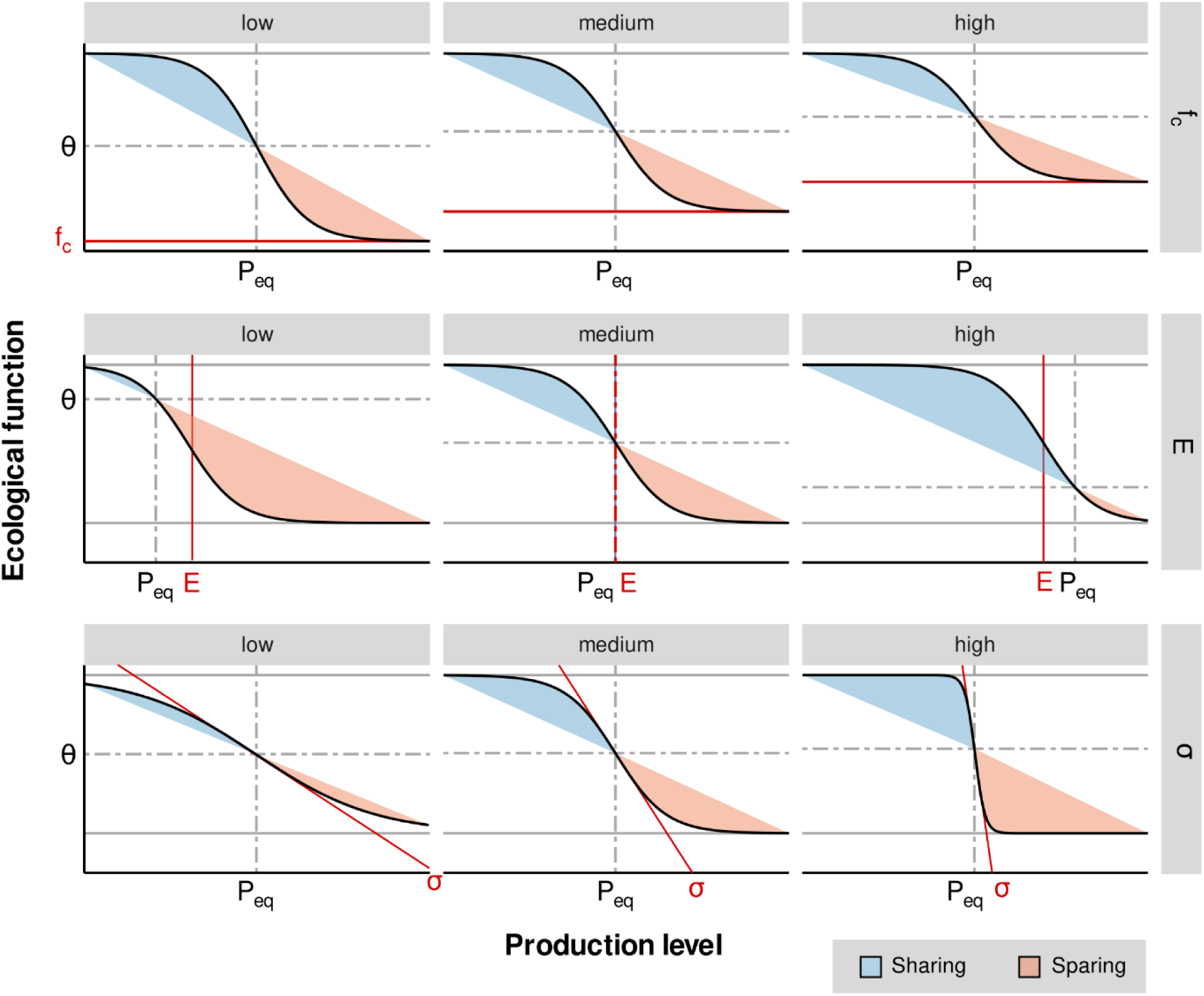
Influence of three parameters of the sigmoidal function on the sharing-sparing balance. P*_eq_* represents the production level and θ the preservation threshold at which the strategies are equivalent.The larger the shaded space between the sinus and the chord, the more advantageous the corresponding strategy.

Sensitivity (*σ)* is closely linked to core function, and less likely to shift the balance of the preferable land use strategy. Nevertheless, high sensitivity and lower core function values can be associated with hysteresis phenomena (Litzow & Hunsicker, 2016), a relevant consideration for non-productive reconversion of cropland. Moreover, low levels of sigma are indicative of small magnitudes in the difference in outcome between the two strategies, approximating what we termed the “ethical” model.

The other two parameters of the sigmoid response model, namely efficiency and preservation threshold, offer more scope for substantial intervention and manipulation. Increasing the preservation threshold, fundamentally a policy intervention, has the effect of excluding scenarios favourable to land sparing, while retaining the land sharing options (Figure 4b).

Enforcing stricter environmental measures is indeed a very effective policy tool if the aim is moving the equilibrium towards low-intensity agriculture (Mupepele et al., 2021; Phalan et al., 2016). However, it involves lowering yields, financial margins and global production compared to restriction-free agricultural systems.

Raising the land sharing peak of the isoerg curve above the land sharing counterpart is the only option to make land sharing more competitive without impacting overall yield. Increased levels of ecological efficiency, i.e., reaching higher yields in the resilience phase of the production curve, can dramatically shift the equilibrium (Figure 6). This would however require a substantial shift of paradigm compared to the recent technological history of agriculture (Evenson & Gollin, 2003). Since the green revolution with the mass use of nitrogen fertiliser produced by the Haber–Bosch process, and advances in crop breeding, global agriculture has witnessed an unprecedented push in the tolerance phase of production systems. In particular, selective breeding for high yielding crops with high chemical and mechanical inputs has given farmers access to the tools for more intensive high-density production. The effect in the sigmoid response framework has been a striking lengthening of the tolerance phase compared to the resilience phase, which resulted in a decline in ecological efficiency at a global scale (i.e., a leftward shift of the transition phase compared to maximum production). This is the ultimate root of the structural disadvantage of land sharing strategies emerging in literature (A. Balmford et al., 2015). Underpinning the tolerance of agricultural systems, attempts at de-intensification are unlikely to be met with success, and their poor performance may discourage further efforts. Identification of less intensive farming strategies will be instrumental in giving land managers the possibility to choose between two competitive strategies for a wider set of environmental conditions and preservation policies, together with large-scale socio-political rethinking (Fischer et al., 2017) and technological advancements.

Concerning the latter, improvements in conservation agriculture techniques and crop diversity with multi-species, and optimised rotations that include cover crops, are important tools to raise the ecological efficiency and yield at low input levels. This would be particularly relevant where soils are degraded after high-intensity production when the stable state is severely depleted compared to the undisturbed state. However, arable techniques alone may only bring marginal improvements, while new crops generated by improved breeding and gene editing could bring dramatic shifts in low input agriculture (Lotz et al., n.d.; Wickson et al., 2016). These developments gave land sparing a competitive advantage across most of the world (B. Balmford et al., 2019). If the competing land-sharing strategy cannot rely on chemical fertilisers, pesticides and aggressive cultivation, a credible path forward would require a vigorous effort towards the targeted selection of plant traits as well as cover and cash crops. These technical improvements would be able to generate an efficient closed system with minimal inputs (Machado et al., 2022). This is especially relevant in the light of global warming and the renewed interest in the development of drought, heat resistant or early-flowering crop varieties (Karavolias et al., 2021).

The limitations of the sigmoid response model and the isoerg and preservation threshold conceptual space are linked to the high-level conceptualisation at its basis. However, they also allow to identify possible opportunities for further research that would benefit from this theoretical foundation and greatly expand its potential. Link functions could be used to map scale effects on spared land or to decouple intensity and agricultural production. Moreover, the addition of landscape-scale production thresholds to preservation targets could be implemented in an integrated ecological-function-to-yield response model.

In summary:

- firstly, de-intensification techniques are predicted to have more chances of succeeding where the loss of ecological function under intensive management is more pronounced.
- second, without enforced protection thresholds and policies for the internalisation of environmental costs, de-intensification techniques are unlikely to be competitive, irrespective of the configuration of the other parameters.
- finally, dramatic improvements in the capability of enhancing production at low chemical and energy inputs, such as targeted genetic improvement of both cover and cash crops to minimise environmental damage and enhance natural biodiversity as well as fine-tuning existing agronomic practices, are the ultimate keys to the global success of alternatives to the prevalent land sparing approach.

Whatever form the evolution in land use patterns will take, the sigmoidal response framework does not only introduce a coherent system in agreement with published literature and experimental results. It also provide insights into the role of de-intensification in the toolkit for sustainable agricultural development of land managers and help predict which interventions can be effective, and in which policy and environmental conditions.

## 8. Author Contributions

MFJ: conceptualization, formal analysis, statistics and simulations, writing and figure generation. NB: conceptualization, writing, mathematical analysis, figure generation. BJR: conceptualization, supervision, writing – review & editing. IS: conceptualization, supervision, writing – review & editing. AJM: conceptualization, funding acquisition, resources, supervision, writing – review & editing. All authors contributed to editing and reviewing.

## 10. Acknowledgments

The authors would like to thank PD Dr Anita Risch for her comments to the draft.

## Contributions to the field

One of the most fiercely disputed topics within the larger context of the ecology versus economy debate revolves around the two competing land use strategies of land sharing and land sparing. Both sides of the ideological divide have theorised models linking ecological function and production. The manuscript offers a conceptual synthesis harmonising apparently contradictory approaches in a coherent framework. It is argued that conflicting views stem from special cases of a more universal function. This reconciliation helps transcend ideological barriers and focus instead on the structural determinants of ecosystem function preservation under productive land use. In addition to providing clarity on the mechanics linking the response of ecological function to increasing levels of land use intensity, the framework potentially allows for predictions to be made about optimal strategies in different environmental contexts and the policy and technological interventions that are necessary for land sharing strategies to have a chance at large scale implementation despite current structural advantages of land sparing. In synthesis, the manuscript provides an innovative conceptual tool to assess optimal strategies for biodiversity preservation and viable agricultural production.

